# Design Principles for Polymerase Strand Recycling Circuits

**DOI:** 10.1101/2025.03.17.643471

**Authors:** Yueyi Li, Arno E. Gundlach, Andrew D. Ellington, Julius B. Lucks

**Affiliations:** Department of Chemical and Biological Engineering, Northwestern University, Evanston, Illinois 60208, USA; Center for Synthetic Biology, Northwestern University, Evanston, Illinois 60208, USA; Department of Molecular Biosciences, University of Texas at Austin, Austin, TX, 78712; Center for Systems and Synthetic Biology, University of Texas at Austin, Austin, TX, 78712; Interdiscipinary Biological Sciences Graduate Program, Northwestern University, Evanston, Illinois 60208, USA; Center for Water Research, Northwestern University, Evanston, Illinois 60208, USA; Center for Engineering Sustainability and Resilience, Northwestern University, Evanston, Illinois 60208, USA

## Abstract

Cell-free biosensing systems are being engineered as versatile and programmable diagnostic technologies. A core component of cell-free biosensors are programmable molecular circuits that improve biosensor speed, sensitivity and specificity by performing molecular computations such as logic evaluation and signal amplification. In previous work, we developed one such circuit system called Polymerase Strand Recycling (PSR) which amplifies cell-free molecular circuits by using T7 RNA polymerase off-target transcription to recycle nucleic acid inputs. We showed that PSR circuits can be configured to detect RNA target inputs as well as be interfaced with allosteric transcription factor-based biosensors to amplify signal and enhance sensitivity. Here we expand the development of PSR circuit design principles to generalize the platform for detecting a diverse set of model microRNA inputs. We show that PSR circuit function can be enhanced through engineering T7 RNAP, and present troubleshooting strategies to optimize PSR circuit performance.

## Introduction

Cell-free biosensing is a rapidly developing technology platform because of its ability to detect a wide range of nucleic acid, protein and chemical targets at the point-of-need without expensive conventional laboratory equipment [1-6]. A key component of cell-free biosensing reactions are programmable molecular circuits that improve biosensor speed, sensitivity and specificity by performing molecular computations such as logic evaluation and signal amplification [7, 8]. Recently there has been an interest in performing these computations with DNA nanotechnology approaches based on programmable nucleic acid interactions [9, 10]. Such DNA computations utilize toehold-mediated strand displacement (TMSD) reactions, a process by which single-stranded DNA molecules can bind to, invade, and displace double stranded DNA ‘gates’ [7], releasing new strands that can participate in downstream interactions. In previous work, we interfaced toehold-mediated strand displacement (TMSD) circuits with allosteric transcription factors-based small molecule biosensors, creating new approaches to multi-input logic evaluation and signal thresholding, and expanding the functionality of cell-free biosensors for example by having them operate with analog-to-digital signal conversion [7]. However, attempts at utilizing TMSD-based catalytic amplification circuits such as catalytic hairpin assembly (CHA) [11], failed due to off-target transcription of DNA gates by T7 RNA polymerase (T7 RNAP) [8, 12, 13].

We recently addressed this limitation through a novel circuit architecture that leverages T7 RNAP off-target transcription to recycle nucleic acid inputs within DNA TMSD circuits called polymerase strand recycling (PSR) [8]. PSR is compatible with T7 RNAP *in vitro* transcription and can detect RNA targets directly as well as enhancing the limit of detection for allosteric transcription factor-based biosensors to detect small molecules [8]. PSR circuit designs are comprised of three main components: RNA input, fuel gate and signal gate (**Figure 1a, Supplementary Figure 1**). Both the fuel gate and signal gate are designed to be double stranded, but leave 5’ and 3’ unpaired overhangs called ‘toeholds’ that facilitate initial binding and strand invasion of the duplex (**Supplementary Figure 1**) [14]. In a PSR reaction, the circuit is activated by RNA inputs, which can be the input signal directly, or can be generated by upstream transcriptional biosensing mechanisms. RNA inputs are converted to DNA strands via strand invading a nucleic acid duplex called a “fuel gate”, which releases a DNA strand called RecycleD. The released RecycleD strand can in turn can invade a “signal gate”, consisting of a DNA duplex containing fluorophore-labeled strand and a quencher-labeled strand, releasing the quencher and generating fluorescent signal. The system is designed such that the resulting RecycleD:fluorophore-strand complex contains a 3’ toehold, which can be transcribed by T7 RNAP, releasing RecycleD for further rounds of signal generation and recycling (**Figure 1a, Supplementary Figure 1**).

**Figure 1.**
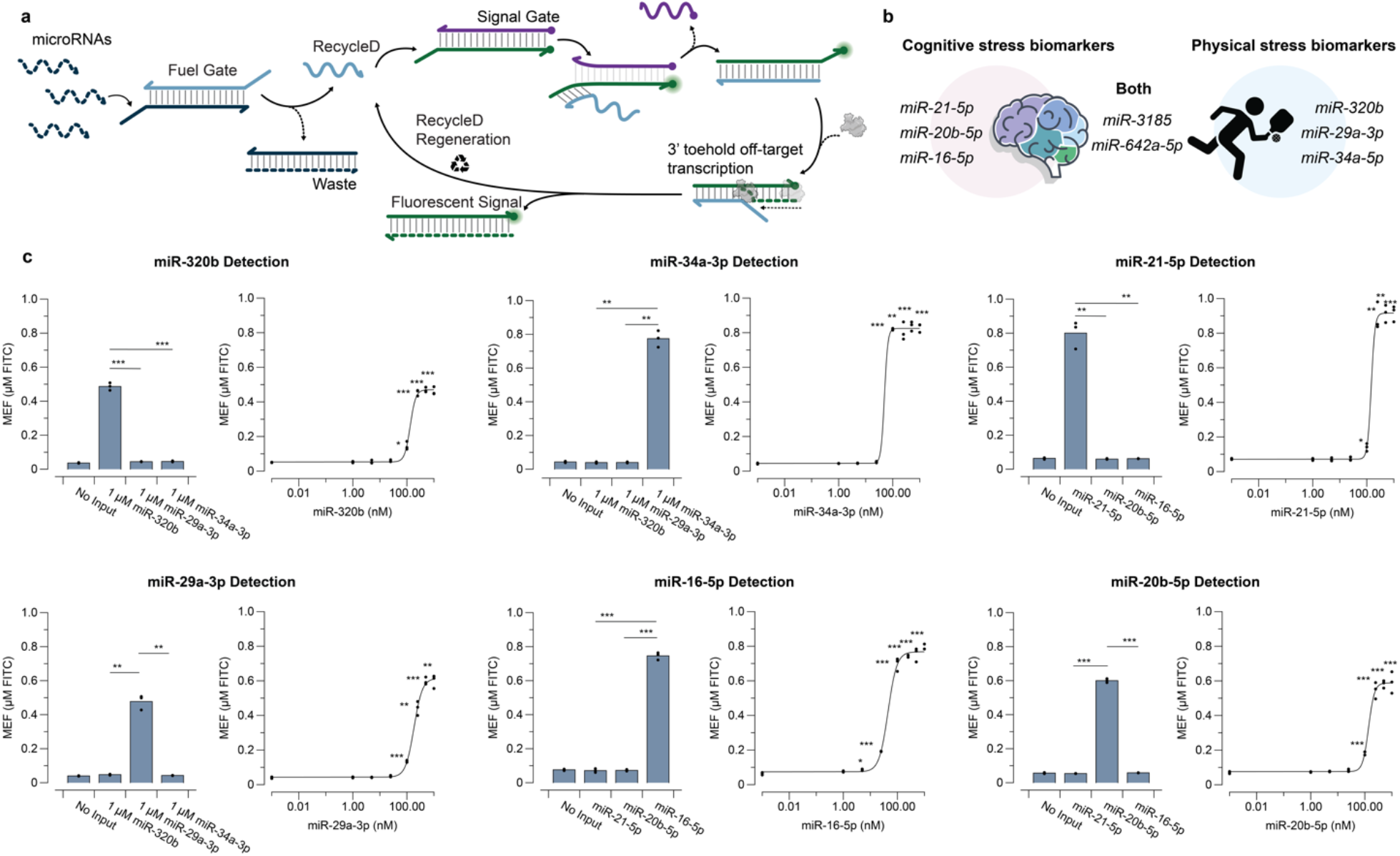
RNA input sensing with PSR. **a.** PSR circuit design for detecting microRNAs. Fuel gates are designed to sequester microRNAs and release RecycleD via toehold-mediated strand displacement. RecycleD is designed to strand invade the signal gate and form a new duplex with the fluorophore-strand, leaving a 4-nt 3’ toehold. T7 RNAP off-target transcription from the 3’ toehold results in RecycleD regeneration and signal amplification. **b**. miR biomarkers that increase or decrease in response to physical stressors, cognitive stressors, or both. **c**. PSR microRNA sensing performance. Different sequences for PSR fuel gates and signal gates were designed to detect purified model miR targets miR-320b, miR-29a-3p, miR-34-3p, miR-16-5p, miR-21-5p, and miR-20b-5p. The PSR circuits containing 1.5 µM of fuel gate, 2.5 µM of signal gate, 2 ng of T7 RNAP all detect 1 µM of their respective miR targets with significant specificity after 2 hrs. Significance values comparing fluorescence from targeted miR inputs and the indicated non-targeted inputs were determined using a two-tailed, paired Student’s t-test. Dose responses with miR-320b, miR-29a-3p, miR-34a-3p, miR-21-5p, miR-20b-5p measured after 2 hrs showed limit of detection (LoD) of 100 nM. Dose response with miR-16-5p measured after 2 hrs shows LoD of 5 nM. The LoDs were determined from the concentration value at which the signal was significantly greater (p < 0.05) than the no input condition using a two-tailed, heteroscedastic Student’s t-test. Data shown in bar graphs in c are n = 3 experimentally independent replicates, each plotted as a point with raw fluorescence standardized to MEF (µM FITC) and bar heights representing the average over these replicates. Data shown in dose response graphs in c are n = 3 experimentally independent replicates, each plotted as a point with raw fluorescence standardized to MEF (µM FITC). Curves in c represent Hill Equation fits (see Methods), with fit parameters in Source Data. The p-value range is indicated by asterisks (***p < 0.001, **p = 0.001–0.01, *p = 0.01–0.05). Exact p-values can be found in Source Data.

In previous work, we demonstrated PSR circuits can directly detect three distinct microRNAs (miRs) with high specificity and limits of detection (LoDs) within 5 – 100 nM range [8]. In this study, we further explored the design principles for PSR circuits by developing seven distinct PSR circuit architectures to detect additional miR inputs. We analyzed the successful and unsuccessful PSR circuit designs across different RNA sequence inputs and developed design principles for optimizing specificity and sensitivity. We also provide evidence that PSR circuit function can be improved through engineering the T7 RNAP enzyme. Overall we demonstrate a general strategy for engineering PSR circuits that could have a range of applications in cell-free biosensing and DNA/RNA nanotechnology and synthetic biology more broadly.

## Results

### PSR circuit architecture can be generalized to detect various RNA inputs

To demonstrate the generalizability of PSR circuits for detecting different RNA input sequences, we chose miRs as model targets. miRs are a class of small, noncoding RNAs that are key regulators of gene expression in eukaryotes, and are important biomarkers of cell-type and health status [15, 16]. Most miRs are approximately 22-nucleotide (nt) long but exhibit a wide range of melting temperatures (T_m_s) due to sequence variation, making them challenging targets for generalizable sensing platforms [17]. Here, we chose six distinct miRs (miR-21-5p, miR-20b-5p, miR-16-5p, miR320b, miR-29a-3p, and miR-34a-5p) that are either upregulated or downregulated from physical stressors, cognitive stressors, or both [18-24] as model inputs for PSR circuits (**Figure 1b**). These miRs have T_m_s ranging from 57ºC – 70ºC and are between 21 – 23-nt in length.

To explore the ability of PSR circuits to detect this range of model input, we first designed modified fuel gate, RecycleD, and signal gate sequences to match the desired miR sequences (**Figure 1a**). We first assessed the specificity of the PSR circuit designs by adding 1 µM of synthesized and purified miR to its respective PSR circuit, as well as two other PSR circuit designs that are intended to detect non-target miRs. Fluorescence characterization demonstrated that all seven PSR designs could only be activated by their target miR targets, each generating fluorescent signals that are at least 9.5-fold higher compared to non-target miRs (**Figure 1c**).

We then assessed the sensitivity of PSR designs by performing a titration with miRs to determine the limits of detection (LoDs), defined as the concentration value at which the signal was significantly greater (p < 0.05) than the no input condition. We determined the LoDs to be within the 5 – 100 nM range for PSR to detect RNA targets. Specifically, miR-320b, miR-29a-3p, miR-34a-3p, miR-21-5p, miR-20b-5p targets showed LoDs of 100 nM, and miR-16-5p target showed LoD of 5 nM (**Figure 1c**).

PSR relies on the function of T7 RNAP to perform the strand recycling reaction (**Figure 1a**). We therefore reasoned that PSR circuit performance could be improved by varying the T7 RNAP itself. We compared wildtype T7 RNAP performance with two variants, M6 and Y639F, to detect miR-29a-3p and found the M6 variant improves fluorescent signal while the Y639F variant hinders fluorescent signal generation compared to wildtype (**Supplementary Figure 2**).

**Figure 2.**
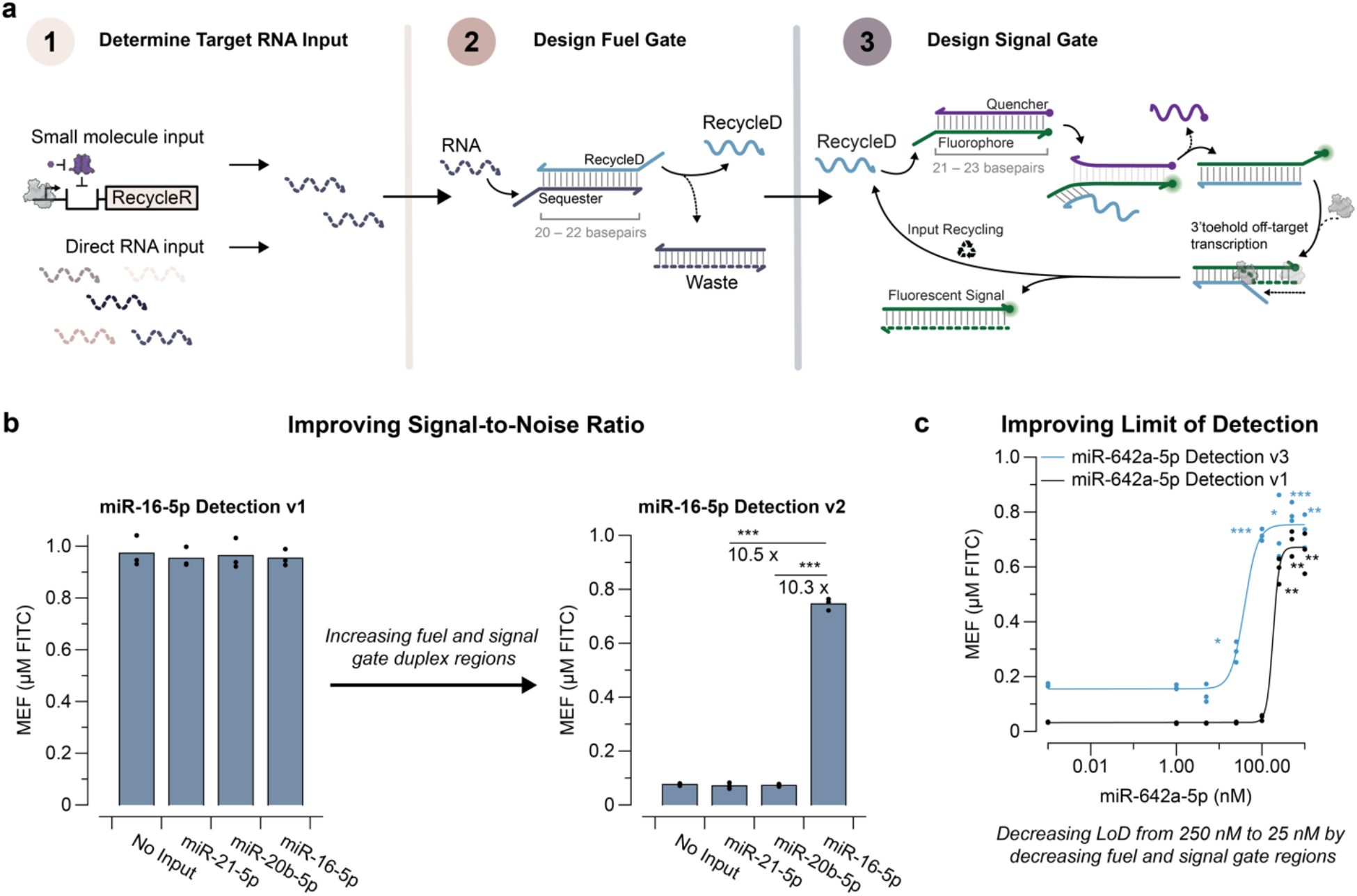
PSR design process and troubleshooting approaches. **a.** Design process for building PSR circuits. (Left) Select RNA inputs used to activate a PSR circuit can either come from transcription from ligand-regulated sensing mechanisms or directly as RNA molecules. (Middle) The PSR fuel gate is formed by two nucleic acid strands with an overlap of 20 – 22-bp, a 5-nt 5’ overhang on RecycleD and a 9 – 10-nt 5’ overhang on the sequester strand. The sequester strand is 2’–O–methyl modified and contains the complementary sequence for the RNA input. RecycleD is a DNA strand that gets released from RNA input strand invasion. (Right) The signal gate is formed by two nucleic acid strands with an overlap of 21 – 23-bp and an 8-nt 5’ overhang on the fluorophore strand. The signal gate is designed to be strand invaded by RecycleD to form a new duplex with the fluorophore-strand containing a 4-nt 3’ toehold to enable T7 RNAP off-target transcription. **b**. Improving signal-to-noise ratio. The PSR circuit designed to detect miR-16-5p containing 1.5 µM of fuel gate, 2.5 µM of signal gate, 2 ng of T7 RNAP initially generated high fluorescent signal in the absence of its target miR after 2 hrs. By increasing the overlapping regions on the fuel and signal gates by 2-bp, the signal-to-noise ratio improved to ∼10-fold (see Supplementary Figure 4). Significance values comparing fluorescence from targeted miR inputs and the indicated non-targeted inputs were determined using a two-tailed, paired Student’s t-test. **c**. Improving LoD for miR detection. The PSR circuit designed to detect miR-642a-5p initially showed an LoD of 250 nM after 2 hrs. By decreasing the overlapping regions on the fuel and signal gates by 2-bp, we demonstrated an LoD of 25 nM (see Supplementary Figure 5). The LoDs were determined from the concentration value at which the signal was significantly greater (p < 0.05) than the no input condition using a two-tailed, heteroscedastic Student’s t-test. The p-value range is indicated by asterisks (***p < 0.001, **p = 0.001–0.01, *p = 0.01–0.05). Exact p-values can be found in Source Data. Data shown in **b** are n = 3 experimentally independent replicates, each plotted as a point with raw fluorescence standardized to MEF (µM FITC) and bar heights representing the average over these replicates. Data shown in c are n = 3 experimentally independent replicates, each plotted as a point with raw fluorescence standardized to MEF (µM FITC). Curves in c represent Hill Equation fits (see Methods), with fit parameters in Source Data.

Overall, these results demonstrated that PSR circuits can be generalized to detect RNA inputs of various sequences and a wide range of T_m_s.

### Establishing design principles and troubleshooting approaches for engineering PSR circuits

Next, we developed design principles that could be applied to achieve the expected performance from PSR circuits (**Figure 2a**), along with troubleshooting approaches that could be used to improve the signal-to-noise ratio or the LoD if the initial circuit design fails to meet performance metrics (**Figure 2b, Figure 2c**).

We developed design principles for fuel and signal gates that could serve as a starting point to design PSR circuits for new RNA input sequences that are miRs or a ∼22-nt region of a longer RNA target. These design principles consist of several features: (1) The fuel gate should be comprised of two nucleic acid strands, sequester and RecycleD with an overlapping region of 20 – 22-basepair (bp). (2) The sequester strand should be 2’–O–methyl modified and contain the complementary sequence for the RNA input, including a 5’ toehold that is 9 – 10-nt in length, as well as a region partially complementary to RecycleD (**Supplementary Figure 1a**). (3) The signal gate should be comprised of a DNA strand with a 3’ fluorophore modification and another DNA strand with a 5’ quencher modification. The gate should have a 21 – 23-bp overlapping region as well as an 8-nt 5’ toehold on the fluorophore-strand (**Supplementary Figure 1b**). (4) The RecycleD:fluorophore-strand complex should contain a 4-nt 3’ toehold on the fluorophore-strand (**Supplementary Figure 1c**). Since the rate of T7 RNAP off-target transcription activity has been found to be dependent on the 3’ toehold sequence [12], we reused the same 4-nt 3’ sequence for all existing PSR circuit designs.

If the initial PSR design did not produce the expected performance, we found that the results could likely be improved with design adjustments to the fuel and signal gates. In several cases we found that initial PSR designs showed high fluorescent leak or LoDs above the expected range of 5 – 100 nM. In these cases, we were able to improve the PSR circuit performance to be in the expected range through adjusting the fuel and signal gate designs (**Supplementary Figure 3–5**). For example, the initial circuit design for miR-16-5p produced high fluorescent signals in the absence of the miR target (**Figure 2b**). By extending RecycleD on the 3’ end by 2-nt, thus increasing the fuel gate overlapping region by 2-bp and subsequentially increasing the signal gate duplex by 2-bp (**Supplementary Figure 3**), the signal-to-noise ratio improved to ∼10-fold (**Figure 2b**). In a previously published PSR circuit to detect miR-642a-5p, we initially demonstrated an LoD of 250 nM, which is above the expected range of 5 – 100 nM (**Figure 2c**). We were able to decrease the LoD to 100 nM by decreasing the overlapping regions on the fuel and signal gate by 1-bp and subsequently to 25 nM by decreasing the overlapping regions on fuel and signal gate by an additional 1-bp (**Figure 2c, Supplementary Figure 5**).

Overall these results demonstrate a general approach for designing and optimizing PSR circuits for RNA detection.

## Discussion

In this study, we expanded the toolbox for building PSR circuits to detect a diverse set of input RNA sequences with greater than 10-fold signal-to-noise ratio and LoD within the range of 5 – 100 nM. We also developed design principles to achieve the expected circuit performance with new target RNA inputs. Even if an initial design fails, the signal-to-noise ratio and LoD can be optimized with adjustments to fuel and signal gates design. In some cases, only the RecycleD or sequester sequence needs to be adjusted to reduce fluorescent signal leak, which is more economic compared to redesigning the entire set of PSR circuit elements (including fluorophore-strand and quencher-strand) (**Supplementary Figure 3)**. If cost-saving is a priority, only extending RecycleD by 1-nt could be a potential initial troubleshooting step.

In previous work, we showed that the major strength of PSR circuit is their ability to interface with transcriptional biosensors to optimize the sensitivity of detection of the large number of allosteric transcription factors (aTFs) used for chemical biosensing applications. aTF-based PSR circuit design is similar to the design principles described above but requires designing a DNA template comprised of the T7 RNAP promoter, a 2-bp spacer, an aTF-operator site, and a target RNA input sequence (RecycleR). To adapt PSR circuit for a new aTF, the corresponding aTF-operator site needs to be inserted between the 2-bp spacer and the RecycleR sequence (**Supplementary Figure 6**). Unless there are downstream sequence constraints, the sequences for RecycleR, fuel and signal gate can be directly adapted from previously published designs. It is interesting to note that the aTF-based PSR circuit fuel and signal gate overlapping regions do not follow the same design rules as purified target RNA detection (**Supplementary Figure 6**). We hypothesize that the overlapping regions could be shorter in aTF-based PSR circuits without significant fluorescent signal leak because there is a T7 RNAP promoter site, which slows down T7 RNAP off-target transcription activities [13].

Because PSR has a unique configuration for signal amplification, and relies on a T7 RNA polymerase mechanism, promoter-less initiation of transcription [25] [7] that is poorly understood, we hypothesized that variants of T7 RNAP might improve circuit performance. To that end, we and others had previously developed a number of T7 RNAP variants with higher thermostabilities and wider substrate specificities [26] [27]. When tested in the context of the PSR circuitry, we observed different T7 variants could either enhance or hinder PSR reactions, suggesting PSR performance could be further tuned and PSR could have additional applications due to the qualities of the variants (**Supplementary Figure 2**).

Other nucleic acid-based miR detection methods, such as catalytic hairpin assembly, have demonstrated high sensitivity in the fM ranges. However, PSR stands out as a detection platform due to its simplicity and programmability. Furthermore, the ability to integrate PSR into transcriptional circuits expands its potential for application in advanced circuit designs.

Overall, we believe that the PSR circuits can serve as a platform technology that can be adapted to detect RNA inputs through PSR gate design, and chemical targets through interfacing with aTF biosensors [8]. We anticipate this toolbox for PSR circuit design will serve important roles in cell-free technologies including in biosensing and further expanding molecular computation in cell-free systems.

## Materials and Methods

### DNA gate preparation

DNA fuel gates and signal gates used in this study were synthesized by Integrated DNA technologies as HPLC or PAGE purified and chemically modified oligos (**Supplementary Data**). Gates were generated by denaturing complementary strands at 95° C for 5 min and slow cooling (–0.1 °C s^−1^) to room temperature in annealing buffer (50 mM Tris-HCl, pH 8.0, 10 mM MgCl_2_). Fuel gates are annealed with 20% excess sequester strands and signal gates are annealed 20% excess quencher-strands. DNA fuel gates and signal gates were stored at 4ºC until use. See Li_Supplementary_Data_1.xlsx for a list of oligo sequences used in this study.

### RNA preparation

Purified miRs, representing processed microRNA sequences, were synthesized and HPLC purified by Integrated DNA Technologies and rehydrated with nuclease-free water. miR aliquots were stored at -80ºC until use. See Li_Supplementary_Data_1.xlsx for a list of miR sequences used in this study.

### PSR reactions

PSR reactions were set up by adding the following components listed at their final concentration: IVT buffer (40 mM Tris–HCl pH 8, 8 mM MgCl_2_, 10 mM dithiothreitol, 20 mM NaCl and 2 mM spermidine), 11.4 mM NTPs pH 7.5, 1.5 µM fuel gate, 2.5 µM signal gate, and MilliQ ultrapure H_2_O to a total volume of 20 µL. Immediately before plate reader measurements, 2 ng of wildtype T7 RNAP and, optionally, purified miRs at the indicated concentration were added to the reaction. T7 RNAP variants were added at a final concentration of 0.2 µM. Reactions were then characterized on a plate reader as described in ‘Plate reader quantification and micromolar equivalent fluorescein standardization’.

### Hill equation fits

Where indicated, data were fit to the Hill equation with the following functional form

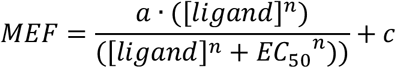

Where [ligand] denotes ligand concentration, *c* represents the response with no ligand, *a* represents the maximum response, and *n* describes the cooperativity. Fitting was performed on individual replicate datasets using DataGraph 5.2 by starting with a = 1.5, EC_50_ = 1, c = 1, n = 1, then optimized for exact parameters with DataGraph 5.2. Curves were generated from fitting the average of the replicates for plotting.

### T7 RNAP variants preparation

Plasmids containing T7 RNAP gene variants with an N-terminal his-tag were transformed into NEB BL21. Colonies were inoculated for overnight culture. Overnight cultures were subcultured the next day 1:100 in superior broth at 37°C. Expression was induced with IPTG at OD .6 and cultured overnight at 18°C. Cells were pelleted through centrifugation at 4,000 g for 10 minutes, resuspended in 30 mL of resuspension buffer (50 mM phosphate buffer [pH 7.5], 300 mM NaCl, 20 mM imidazole, 0.1% Igepal CO-630, 5 mM MgSO4), and lysed through sonication. The lysate was centrifuged at 35,000 g for 30 minutes. A gravity column was assembled with 1 mL of Ni-NTA resin and equilibrated with 10 mL equilibration buffer (50 mM phosphate buffer [pH 7.5], 300 mM NaCl, 20 mM imidazole). The supernatant was applied to a gravity column containing Ni-NTA resin, washed with 20 mL equilibration buffer, then washed with 5 mL wash buffer (50 mM phosphate buffer [pH 7.5], 300 mM NaCl, 50 mM imidazole), and eluted with 3 mL elution buffer (50 mM phosphate buffer [pH 7.5], 300 mM NaCl, 250 mM imidazole). The eluate was dialyzed twice in 2 L Ni-NTA buffer (40 mM Tris-HCl [pH 7.5], 100 mM NaCl, 1 mM DTT, 0.1% Igepal CO-630), once for 4 hours, then overnight. A final dialysis was performed with 1 L buffer (50% Glycerol, 50 mM Tris-HCl [pH 8.0], 50 mM KCl, 0.1% Tween-20, 0.1% Igepal CO-630). The enzyme concentration was measured using the Bradford assay and diluted using the final dialysis buffer. Protein samples were stored at -20°C. until use.

### Plate reader quantification and micromolar equivalent fluorescein standardization

A National Institute of Standards and Technology traceable standard (Invitrogen, catalog no. F36915) was used to convert arbitrary fluorescence measurements to micromolar equivalent fluorescein (MEF). Serial dilutions from a 50 µM stock were prepared in 100 mM sodium borate buffer at pH 9.5, including a 100 mM sodium borate buffer blank (total of 12 samples). For each concentration, three replicates of samples were created in batches of three, and fluorescence values were read at an excitation wavelength of 490 nm and emission wavelength of 525 nm for 6-FAM (fluorescein)-activated fluorescence (Synergy H1, BioTek Gen5 v.2.04). Fluorescence values for a fluorescein concentration in which a single replicate saturated the plate reader were excluded from the analysis. The remaining replicates (three per sample) were then averaged at each fluorescein concentration, and the average fluorescence value of the blank was subtracted from all values. Linear regression was then performed for concentrations within the linear range of fluorescence (0–3.125 µM fluorescein) between the measured fluorescence values in arbitrary units and the concentration of fluorescein to identify the conversion factor. For each plate reader, excitation, emission and gain setting, we found a linear conversion factor (setting the y intercept to 0) that was used to correlate arbitrary fluorescence values to MEF (**Supplementary Figure 7**).

For reaction characterization, 19 µL of reactions were loaded onto a 384-well optically clear, flat-bottom plate using a multichannel pipette, covered with a plate seal and measured on a plate reader (Synergy H1, BioTek Gen5 v.2.04). Kinetic analysis of 6-FAM (fluorescein)-activated fluorescence was performed by reading the plate at 1-min intervals with excitation and emission wavelengths of 490 and 525 nm for 2 hrs at 37 °C. Arbitrary fluorescence values were then converted to MEF by dividing by the appropriate calibration conversion factor.

## STATISTICS AND REPRODUCIBILITY

The number of replicates and types of replicates performed are described in the legend of each figure. Individual data points are shown, and where relevant, the average ± s.d. is shown; this information is provided in each figure legend. The type of statistical analysis performed in Figures 1, 2, SI Figure 2, 3, 4 is described in the legend to each figure. Exact p-values along with degrees from the statistical analysis can be found in Source Data.

## Supporting information

All source data

## DATA AVAILABILITY

All data presented in this paper are available as Source Data and as Supplementary Data. Source Data are provided with this paper.

## AUTHOR CONTRIBUTIONS

Y.L. and J.B.L. designed the study. Y.L. and J.B.L. analyzed the data. Y.L. and A. E. G. conducted the research. Y.L. and J.B.L. developed the methodology. Y.L. and J.B.L. undertook visualization of the data. Y.L. and J.B.L. curated the data. J.B.L. and A.D.E. acquired funding for the study. J.B.L. managed and coordinated the study. J.B.L. and A.D.E. supervised the research.

## ACKNOWLEDEGMENTS

We thank A. Moreno (Northwestern University) for managing the experimental reagents and equipment used in this study; S. O. Kelley (Northwestern University), E.H. Sargent (Northwestern University), M. C. Jewett (Stanford University), and J. L. Chavez (AFRL) for helpful discussions on miR target selection. Y.L. was supported by the National Institutes of Health Training Grant (T32GM008449) through Northwestern University’s Biotechnology Training Program and the Ryan Fellowship and the International Institute for Nanotechnology at Northwestern University. This work was also supported by Army Contracting Command (W52P1J-21-9-3023) and the Defense Advanced Research Projects Agency (DARPA) (N660012324041) to A.D. E. and J.B.L., and the National Science Foundation (2310382) to J.B.L. A.E.G. was supported by the Defense Advanced Research Projects Agency (DARPA) (N660012324041) and by the National Science Foundation (2419640); A.D.E was supported by the Welch Foundation (F-1654). The views, opinions, and/or findings expressed are those of the authors and should not be interpreted as representing the official views or policies of the Department of Defense, the National Science Foundation or the U.S. Government.

