## Supplementary Information for "Design Principles for Polymerase Strand Recycling Circuits"

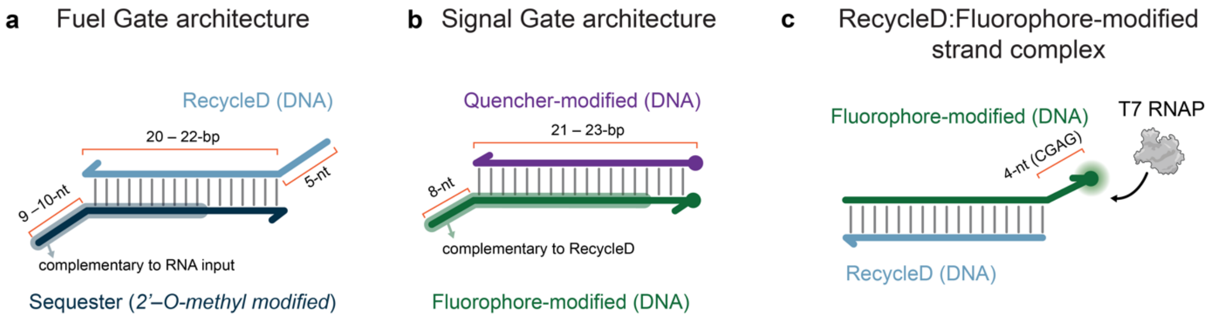


**Supplementary Figure 1 | Design elements in PSR circuits for RNA detection. a**. Fuel gate design architecture for RNA detection. The fuel gate is designed with the sequester strand complementary to the target RNA. Invasion of the fuel gate by the target RNA releases RecycleD. 2'-O-methyl modifications of the sequester strand stabilizes the fuel gate and reduces leak. **B.** Signal gate design architecture for RNA detection. The signal gate is designed to be invaded by RecycleD, releasing the quencher-modified strand and creating a RecycleD:Fluorophore-modified strand complex that fluoresces. **C.** RecycleD:Fluorophore-modified strand complex. The complex has a 4-nt 3' toehold on the fluorophore-modified strand to enable T7 RNAP off-target transcription of this strand, which releases RecycleD.

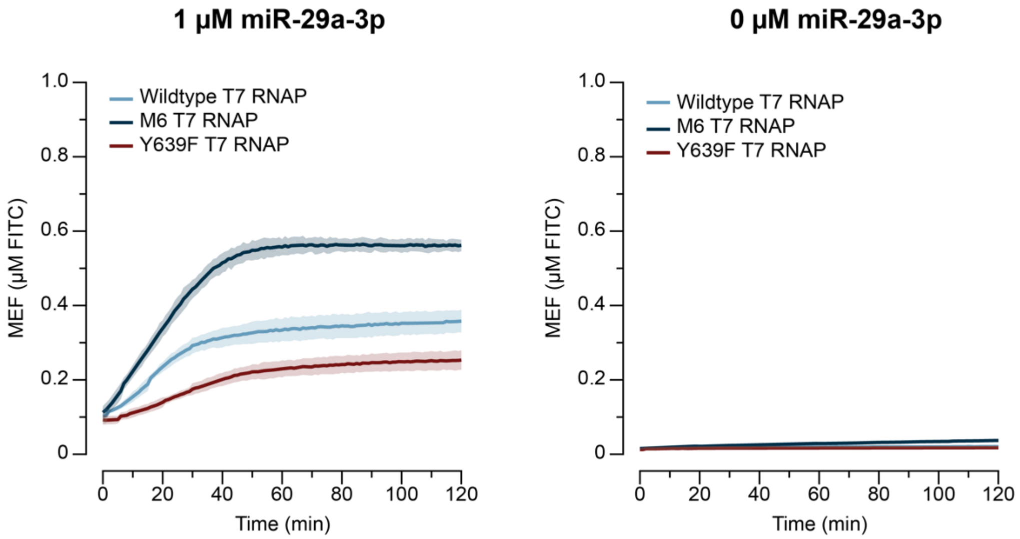


**Supplementary Figure 2 | T7 RNAP variants impact PSR circuit performance.** The kinetic traces of the PSR circuit containing 0.2 μM of either wildtype T7 RNAP, M6 T7 RNAP and Y639F T7 RNAP, 1.5 μM of fuel gate, and 2.5 μM of signal gate with (left) and without (right) 1 μM of added target miR, showing that different T7 RNAP variants could either improve or hinder PSR circuit performance. The data shown are n = 3 independent biological replicates for each condition, with the lines representing the average of the replicates. The shading indicates the average of the replicates ± standard deviation.

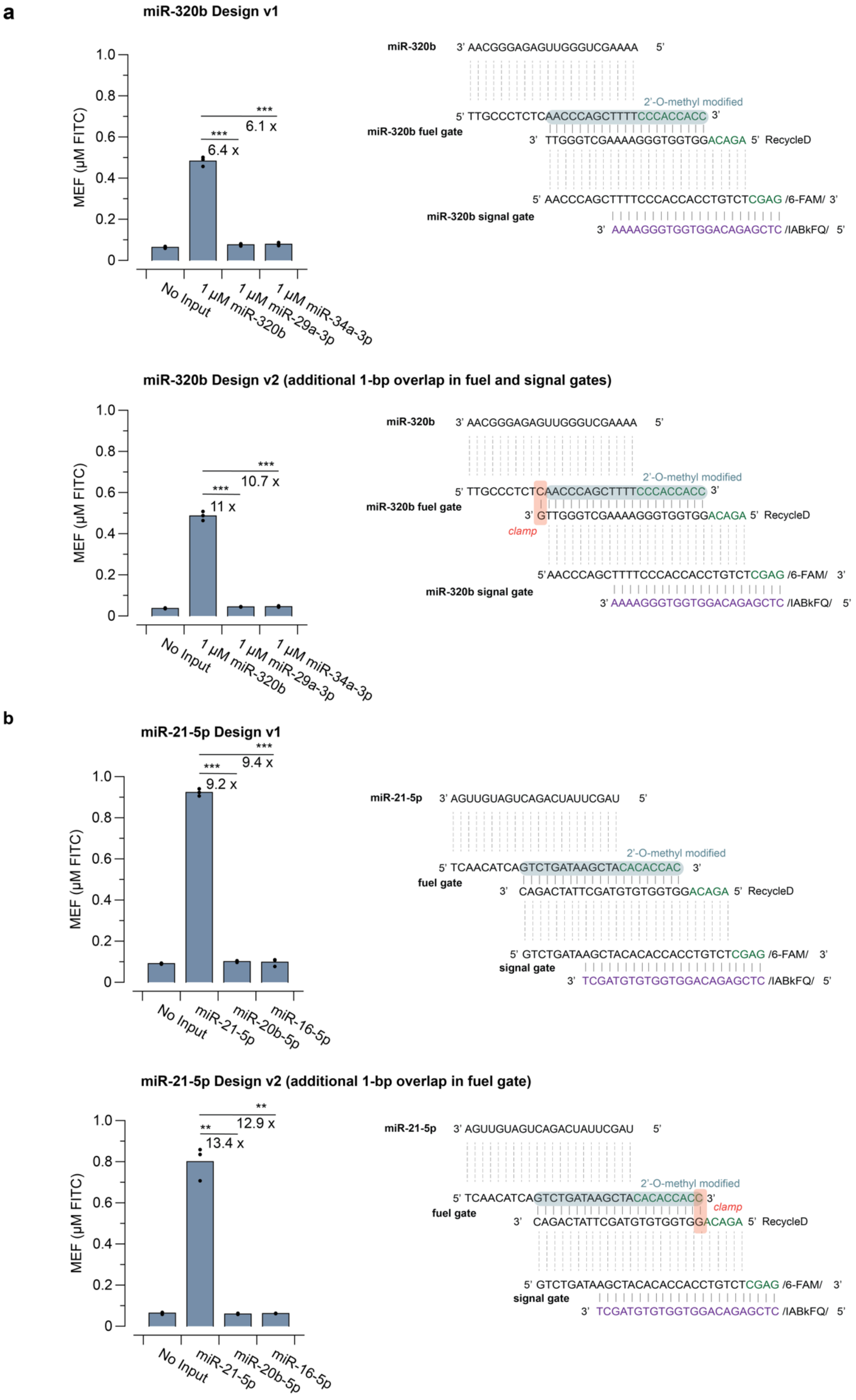
a
miR-320b Design v1
MEF (µM FITC)
1.0
0.8
0.6
0.4
0.2
0
***
6.1 x
***
6.4 x
No Input
1 µM miR-320b
1 µM miR-29a-3p
1 µM miR-34a-3p
miR-320b
3' AACGGGAGAGUUGGGUCGAAAA 5'
2'-O-methyl modified
5' TTGCCCTCTCAACCCAGCTTTTCCCACCACC 3'
miR-320b fuel gate
3' TTGGGTCGAAAAGGGTGGTGGACAGA 5' RecycleD
5' AACCCAGCTTTTCCCACCACCTGTCTCGAG /6-FAM/ 3'
miR-320b signal gate
3' AAAAGGGTGGTGGACAGAGCTC /IABkFQ/ 5'
miR-320b Design v2 (additional 1-bp overlap in fuel and signal gates)
***
10.7 x
***
11 x
clamp
5' TTGCCCTCTCAACCCAGCTTTTCCCACCACC 3'
3' GTTGGGTCGAAAAGGGTGGTGGACAGA 5' RecycleD
5' AACCCAGCTTTTCCCACCACCTGTCTCGAG /6-FAM/ 3'
3' AAAAGGGTGGTGGACAGAGCTC /IABkFQ/ 5'
b
miR-21-5p Design v1
***
9.4 x
***
9.2 x
No Input
miR-21-5p
miR-20b-5p
miR-16-5p
miR-21-5p
3' AGUUGUAGUCAGACUAUUCGAU 5'
2'-O-methyl modified
5' TCAACATCAGTCTGATAAGCTACACACCAC 3'
fuel gate
3' CAGACTATTCGATGTGTGGTGGACAGA 5' RecycleD
5' GTCTGATAAGCTACACACCACCTGTCTCGAG /6-FAM/ 3'
signal gate
3' TCGATGTGTGGTGGACAGAGCTC /IABkFQ/ 5'
miR-21-5p Design v2 (additional 1-bp overlap in fuel gate)
**
12.9 x
**
13.4 x
5' TCAACATCAGTCTGATAAGCTACACACCACC 3'
clamp
3' CAGACTATTCGATGTGTGGTGGACAGA 5' RecycleD
5' GTCTGATAAGCTACACACCACCTGTCTCGAG /6-FAM/ 3'
3' TCGATGTGTGGTGGACAGAGCTC /IABkFQ/ 5'

**Supplementary Figure 3 | Increasing fuel gate base pairs improves signal-to-noise ratio. a.** miR-320b detection PSR circuit performance improvement. Extending RecycleD by 1-nt, thus introducing a 1-bp clamp on the fuel gate, improves the signal-to-noise ratio in detecting miR-320b compared to off-target miRNA sequences from 6.1–6.4x to 10.7-11x. **b.** miR-21-5p detection PSR circuit performance improvement. Introducing a 1-bp clamp on fuel gate by extending the sequester strand improves the signal-to-noise ratio in detecting miR-21-5p compared to off-target miRNA sequences from 9.2–9.4x to 12.9–13.4x. Data shown in a and b are n = 3 experimentally independent replicates from PSR reactions containing 1.5 μM of fuel gate, 2.5 μM of signal gate, and 2 ng of T7 RNAP, with fluorescence measured after 2 hrs. Each replicate is plotted as a point with raw fluorescence standardized to MEF (μM FITC) with bar heights representing the average over these replicates. Significance values comparing fluorescence from targeted miRNA inputs and the indicated non-targeted inputs were determined using a two-tailed, paired Student's t-test. The p-value range is indicated by asterisks (***$p < 0.001$, **$p = 0.001$–$0.01$, *$p = 0.01$–$0.05$). Exact p-values can be found in Source Data.

**miR-16-5p Design v1**

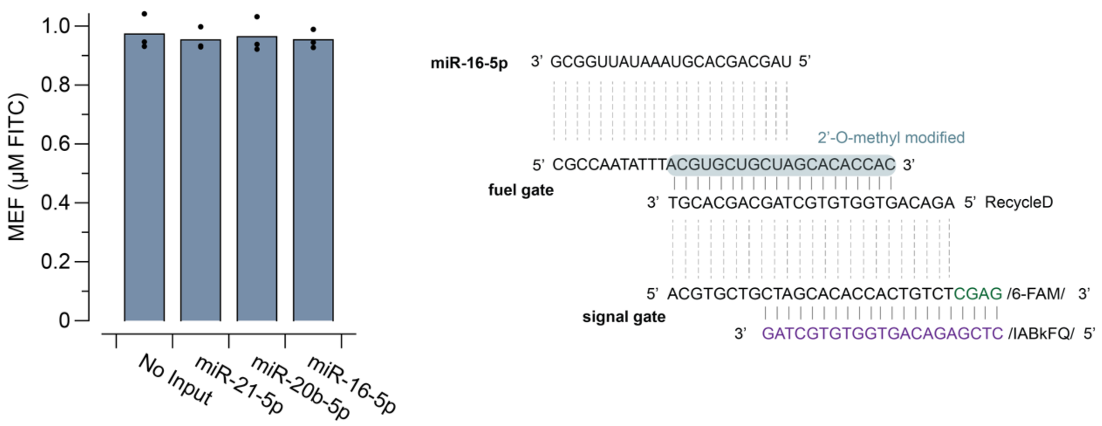


**miR-16-5p Design v2**

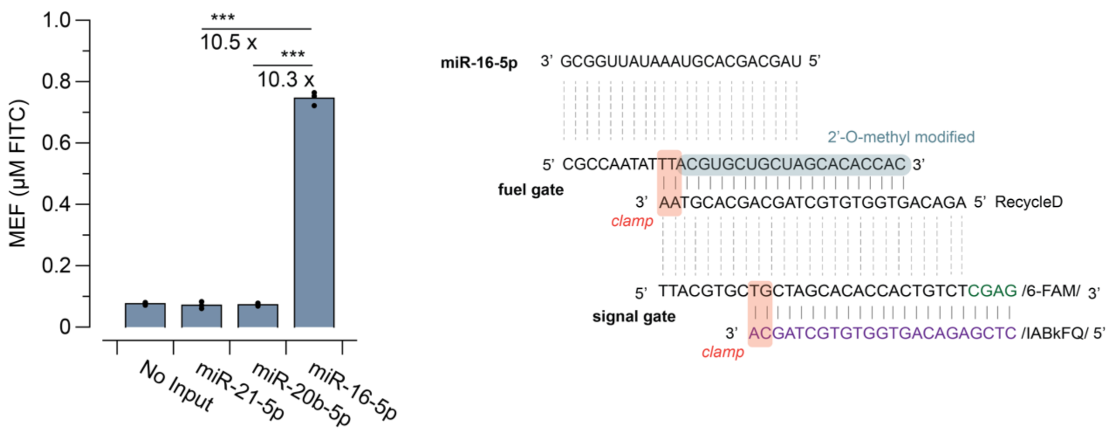


**Supplementary Figure 4 | Increasing fuel and signal gate overlapping regions fixes broken-on PSR circuit for miR-16-5p detection.** Introducing a 2-bp clamp on both the fuel and signal gates improves the signal-to-noise ratio from a nonfunctional, broken ON, PSR circuit (top), to a PSR circuit that shows 10.3–10.5x fold change in detecting miR-16-5p compared to off-target miRNA sequences (bottom). Data shown are n = 3 experimentally independent replicates from PSR reactions containing 1.5 µM of fuel gate, 2.5 µM of signal gate, and 2 ng of T7 RNAP, with fluorescence measured after 2 hrs. Each replicate is plotted as a point with raw fluorescence standardized to MEF (µM FITC) with bar heights representing the average over these replicates. Significance values comparing fluorescence from targeted miRNA inputs and the indicated non-targeted inputs were determined using a two-tailed, paired Student's t-test. The p-value range is indicated by asterisks (***$p < 0.001$, **$p = 0.001–0.01$, *$p = 0.01–0.05$). Exact p-values can be found in Source Data.

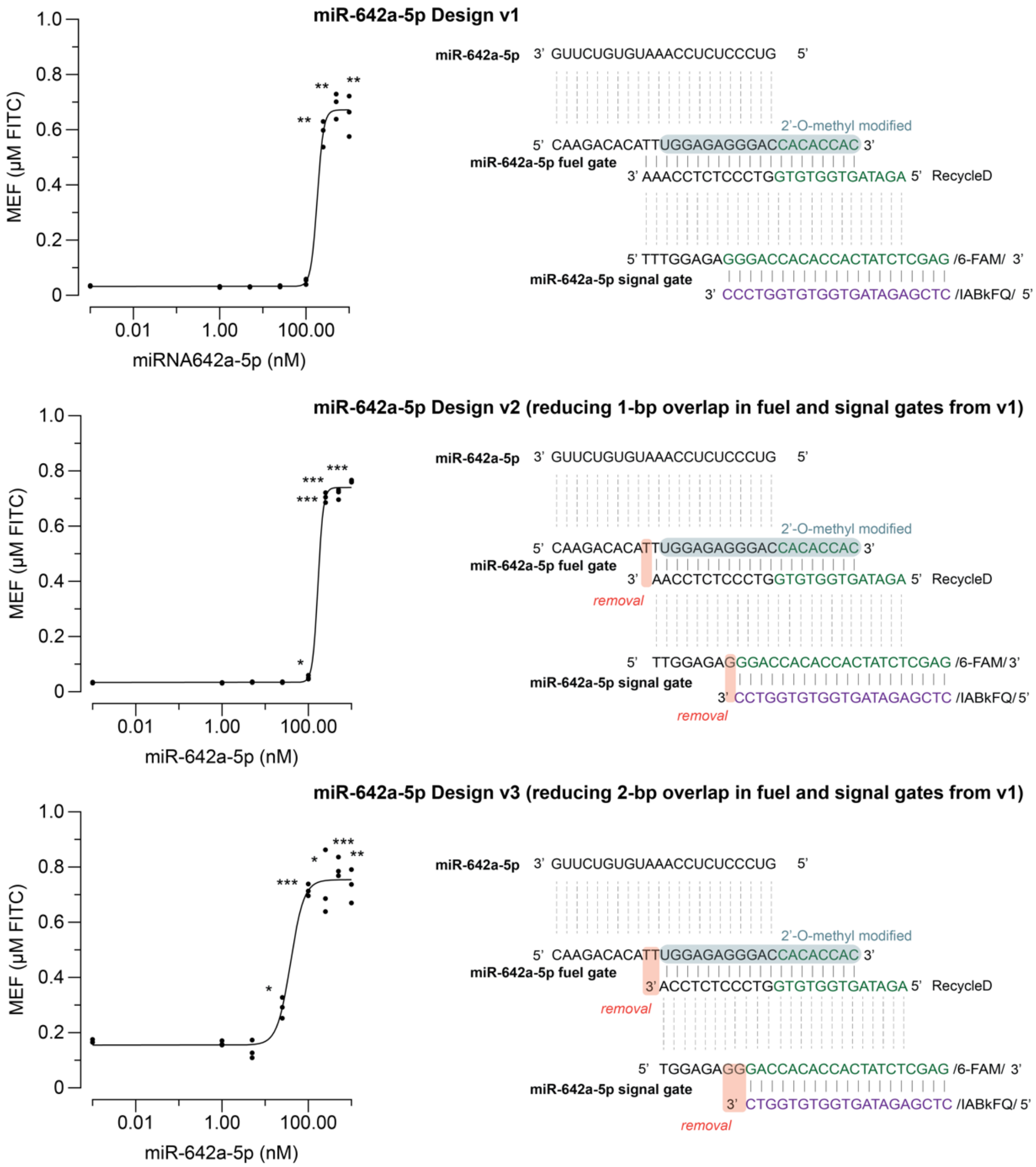


**Supplementary Figure 5 | Decreasing overlapping regions on fuel and signal gates increases LoD for miR detection using PSR.** Different designs of PSR circuits containing 1.5 µM of fuel gate, 2.5 µM of signal gate, and 2 ng of T7 RNAP were used to characterize dose responses to detecting miR-642a-5p measured after 2 hrs. The LoD decreased successively from 250 nM (top) to 100 nM (middle) and 25 nM (bottom) by reducing the overlapping regions on fuel and signal gates by 1–2-bp. The LoDs were determined from the concentration value at which the signal was significantly greater ($p < 0.05$) than the no input condition using a two-tailed, heteroscedastic Student's t-test. The p-value range is indicated by asterisks (***$p < 0.001$, **$p = 0.001–0.01$, *$p = 0.01–0.05$). Exact p-values can be found in Source Data. Data shown are n = 3 experimentally independent replicates, each plotted as a point with raw fluorescence standardized to MEF (µM FITC). Curves represent Hill Equation fits (see Methods), with fit parameters in Source Data.

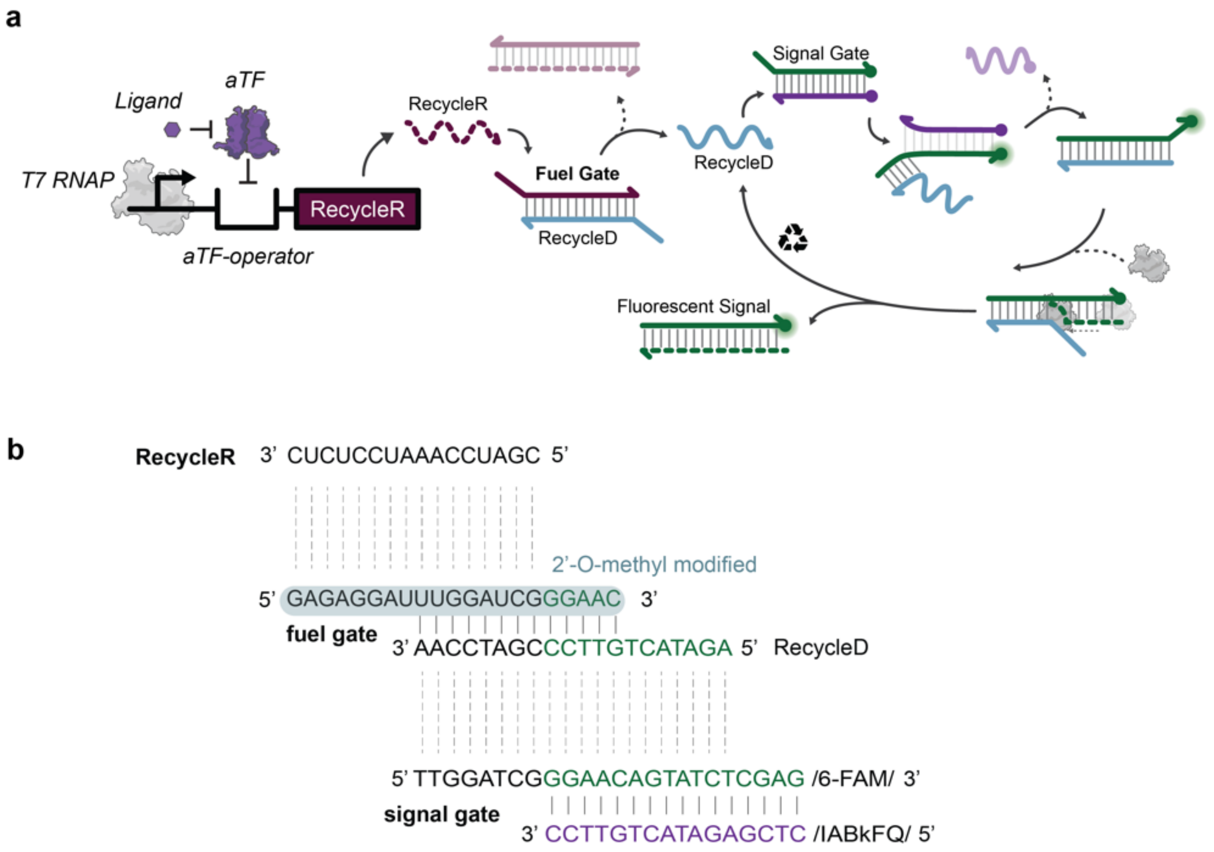


**Supplementary Figure 6. aTF-based biosensing with PSR. a.** PSR circuit scheme for detecting small molecules. RecycleR transcription can be allosterically regulated with a DNA template configured to bind a purified aTF via an operator sequence placed downstream of the T7 promoter and upstream of the RecycleR coding sequence. Activation of the aTF biosensor in the presence of a target ligand generates a signal which is amplified by the downstream PSR circuit. **b.** Design elements in an aTF-based PSR circuit. Sequences for RecycleR, fuel gate, and signal gate. The fuel gate can be strand invaded by RecycleR. It consists of a 2'-O-methyl modified strand and a DNA strand, RecycleD. The DNA signal gate has a fluorophore-labeled strand and a quencher-labeled strand. (Reproduced from Li, Y., Lucci, T., Villarruel Dujovne, M. et al. A cell-free biosensor signal amplification circuit with polymerase strand recycling. Nat Chem Biol (2025). https://doi.org/10.1038/s41589-024-01816-w)

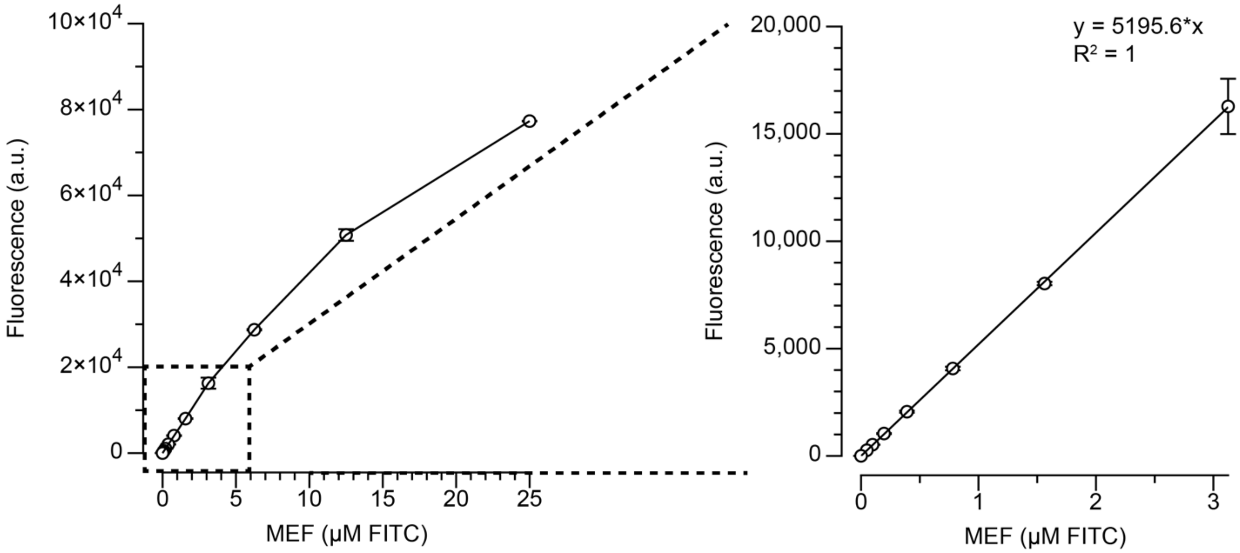


**Supplementary Figure 7. Micromolar Equivalent Fluorescein (MEF) standardization.** Arbitrary units of fluorescence were standardized to μM equivalent fluorescein (μM FITC) using a NIST traceable standard (see Methods). In the representative example shown here, a dilution series of FITC standard was prepared in buffer (100 mM sodium borate, pH 9.5) and measured on a plate reader using the same settings for measuring 6-FAM signal (490 nm excitation, 525 nm emission). The resulting curve, calculated over the linear range of 0–3.125 μM, was then used to standardize fluorescence measured from PSR reactions. The standard curve was generated at regular intervals for each plate reader and each measurement setting. Data shown are for n=3 experimentally independent replicates for each concentration. Error bars indicate standard deviation computed over n=3 replicates.

**Supplementary Files Descriptions**

**Li_Supplementary_Data_1.xls**

DNA, RNA and protein sequences used in this study.

**Li_Supplementary_Data_2.xlsx**

Excel worksheet describing assembly of PSR reactions.
